# Aerosol Formation During Processing of Potentially Infectious Samples on Roche Immunochemistry Analyzers (cobas e analyzers) and in an End-to-End Laboratory Workflow to Model SARS-CoV-2 Infection Risk for Laboratory Operators

**DOI:** 10.1101/2022.02.08.479519

**Authors:** Géza V. Burghardt, Markus Eckl, Doris Huether, Oliver H.D. Larbolette, Alessia Lo Faso, Beatus R. Ofenloch-Haehnle, Marlene A. Riesch, Rolf A. Herb

## Abstract

**Background:** This study assessed formation of potentially infectious aerosols during processing of infectious samples in a real-world laboratory setting, which could then be applied in the context of severe acute respiratory syndrome coronavirus 2 (SARS-CoV-2).

**Methods:** This two-part study assessed aerosol formation when using cobas e analyzers only and in an end-to-end laboratory workflow. To estimate aerosol formation, recombinant hepatitis B surface antigen (HBsAg) was used as a surrogate marker for infectious virus particles to evaluate the potential risk of SARS-CoV-2 infection to laboratory operators. Using the HBsAg model, air sampling was performed at different positions around the cobas e analyzers and in four scenarios reflecting critical handling and/or transport locations in an end-to-end laboratory workflow. Aerosol formation of HBsAg was quantified using the Elecsys^®^ HBsAg II quant II assay. The model was then applied to a SARS-CoV-2 context using SARS-CoV-2 infection-specific parameters to calculate viral RNA copies.

**Results:** Following application to SARS-CoV-2, the mean HBsAg uptake per hour when recalculated into viral RNA copies was 1.9 viral RNA copies across the cobas e analyzers and 0.87 viral RNA copies across all tested scenarios in an end-to-end laboratory workflow. This corresponds to a maximum aspiration rate of <16 viral RNA copies during an 8-hour shift when using cobas e analyzers and/or in an end-to-end laboratory workflow.

**Conclusions:** The low production of marker-containing aerosol when using cobas e analyzers and in an end-to-end laboratory workflow is consistent with a remote risk of laboratory-acquired SARS-CoV-2 infection for laboratory operators.

**Summary:** This study investigated the formation of potentially infectious aerosols during processing of infectious samples in a model using hepatitis B surface antigen (HBsAg) as a marker for infectious virus particles. The risk to laboratory operators of severe acute respiratory syndrome coronavirus 2 (SARS-CoV-2) infection was then inferred. Air sampling was performed around cobas e analyzers and in an end-to-end laboratory workflow, after which HBsAg was quantified and applied to SARS-CoV-2 using SARS-CoV-2 infection-specific parameters. The maximum aspiration rate of <16 viral RNA copies/8-hour shift, when applied to a SARS-CoV-2 context, poses a remote risk of SARS-CoV-2 infection to laboratory operators.

## Introduction

Coronavirus disease 2019 (COVID-19), which was declared a worldwide pandemic in March 2020 (1), is caused by severe acute respiratory syndrome coronavirus 2 (SARS-CoV-2) (2). Widespread testing for SARS-CoV-2 helps prevent the spread of disease by identifying people in the early stages of infection, facilitating disease surveillance and thus a targeted approach for dealing with any outbreaks, and ensures appropriate, timely treatment for patients if required (3, 4). This highlights the importance of laboratory operators carrying out tests for SARS-CoV-2 as key workers in the context of the current pandemic; however, operators could be at a heightened risk of SARS-CoV-2 infection due to exposure to potentially infectious specimens (5).

Aerosols and droplets are the main forms of transmission of SARS-CoV-2 (6); therefore, all laboratory practices should be performed in a way to minimize aerosol and droplet formation to reduce the risk of SARS-CoV-2 infection to laboratory operators (7-9). The processing of samples for SARS-CoV-2 testing by laboratory operators could result in the formation of aerosols, which, if the samples are positive, represents a potentially increased risk of SARS-CoV-2 infection to laboratory operators. There is no clear understanding of infection risk in the laboratory setting (10) and aerosol formation during processing of potentially infectious samples within a SARS-CoV-2 context has yet to be investigated.

The Elecsys^®^ SARS-CoV-2 Antigen test (Roche Diagnostics International, Rotkreuz, Switzerland) is an electrochemiluminescence immunoassay used for in vitro qualitative detection of the SARS-CoV-2 nucleocapsid protein in nasopharyngeal and oropharyngeal swab samples. The procedure involves the processing of potentially infectious SARS-CoV-2 samples on cobas e analyzers (Roche Diagnostics International), often as part of an automated end-to-end laboratory workflow. The end-to-end laboratory workflow involves key processes of pre-analytical sample preparation, sample transport to analyzers and automated output or archive solutions, where laboratory operators could be at risk of SARS-CoV-2 infection (10).

One challenge of investigating aerosol formation in a SARS-CoV-2 context is identifying a surrogate marker for SARS-CoV-2-positive samples. Recombinant hepatitis B surface antigen (HBsAg) was deemed a suitable surrogate for use in this study as it self-assembles to form virus-like particles that are assumed to distribute via aerosols, which allows comparison with typical airborne infectious agents (11, 12). Recombinant HBsAg is also comparable, in order of magnitude, to the size of the SARS-CoV-2 virus particle (11, 13). Furthermore, highly positive non-infectious HBsAg samples are stable over time and HBsAg can be detected with very high sensitivity over a wide dynamic range using the Elecsys HBsAg II quant II assay (Roche Diagnostics International).

The aim of this study was to assess formation of potentially infectious aerosols when processing infectious samples using an HBsAg model on a panel of cobas e analyzers and in an automated end-to-end laboratory workflow, similar to the conditions in which the Elecsys SARS-CoV-2 Antigen test would be run. The model was then used to assess the risk to laboratory operators of SARS-CoV-2 infection due to the formation of contaminated aerosols.

## Materials and Methods

This study comprised two sub-studies, the first of which assessed aerosol formation when handling potentially infectious samples using a panel of cobas e analyzers (conducted at Roche Diagnostics, Penzberg, Germany). The second sub-study assessed aerosol formation when handling potentially infectious samples using cobas e analyzers in an end-to-end laboratory workflow (conducted at Roche Diagnostics International, Rotkreuz, Switzerland).

### THE HBsAg MODEL SYSTEM

Recombinant HBsAg (Roche Diagnostics International) was used as a marker for aerosol formation. Two different batches of HBsAg were prepared for the cobas e analyzer panel sub-study and for the end-to-end laboratory workflow sub-study. In an area with a high prevalence of SARS-CoV-2 infection, it would be expected that 10% of samples submitted to a clinical laboratory would be positive and 1% would be highly positive. In the cobas e analyzer panel sub-study, 10% of samples contained HBsAg marker, which simulated a worst-case scenario. In the end-to-end laboratory workflow sub-study, only samples containing HBsAg were used. The highly positive HBsAg samples had a concentration of 1.2–2.5 ×10^6^ IU/mL (determined by measurements of dilution series across the panel of cobas e analyzers) in the cobas e analyzer panel sub-study and a concentration of approximately 1×10^6^ IU/mL (determined by measurement of a dilution series on a cobas e 801 analytical unit) in the end-to-end laboratory workflow sub-study. For extraction purposes, negative and positive samples contained an equivalent volume of HBsAg-specific diluent ([HBSAGQ2 Dil HepB] from the Elecsys HBsAg II quant II immunoassay kit (Roche Diagnostics International) and were used to exclude unknown contamination within the experimental setup and laboratory environment. The negative sample in Elecsys SARS-CoV-2 assay runs was viral transport medium (final concentration: 2% fetal bovine serum, 100 μg/mL gentamycin and 0.5 μg/mL amphotericin B in sterile Hanks balanced salt solution 1X with calcium and magnesium ions) produced at Roche Diagnostics, Penzberg for internal use only. To establish a baseline for HBsAg measurements, blank filters were eluted and then quantified with the Elecsys HBsAg II quant II assay.

### AEROSOL FORMATION DURING USE OF COBAS E ANALYZERS

Prior to the main experiments, two air sampling plausibility checks (details provided in the Supplemental Methods; Supplemental Fig. 1) were carried out in an open, aerosol-allowing environment. This was for the purposes of demonstrating that (i) intense handling of HBsAg-positive samples can produce HBsAg aerosol particles and (ii) that the air sampling technique used in all experiments was able to capture HBsAg aerosol particles, including the elution of HBsAg with HBsAg-specific diluent (HBSAGQ2 Dil HepB) from the filters and quantification with the Elecsys HBsAg II quant II assay.

Aerosol formation was investigated on the cobas e 801 analytical unit (cobas^®^ 8000 configuration), cobas e 402 analytical unit, cobas e 601 module, cobas e 411 analyzer and the cobas e 801 in combination with cobas pro integrated solution (quattro-BB-configuration, equipped with four cobas e 801 analytical units) analyzers; these units were selected because they all include testing modules for respiratory pathogens. Samples were processed using the pipetting sequences and application of the Elecsys SARS-CoV-2 Antigen test. All cobas e analyzers were operated under routine laboratory conditions.

Air sampling was conducted using Gilian GilAir Plus Air Sampling Pumps (Sensidyne, St. Petersburg, FL) equipped with polycarbonate filters (0.8 μm pore size; SKC Inc., Omega Division, Eighty Four, PA). The pumps were operated at a sampling air stream of typically 2000 mL/min according to manufacturer’s instructions for 4.5–8.3 hours, in combination with the IOM sampler (SKC Inc.), and were calibrated at the beginning of each day with the Mesalabs Defender 520-M (Mesa Laboratories Lakewood, Lakewood, CO). The air sampling devices were placed around the fan outlets of the cobas e analyzers at shoulder/head height of the laboratory operator and at further distances from the cobas e analyzers through placement around the different laboratory rooms and computer workstations. Further details of the experimental conditions for air sampling around the cobas e analyzers are provided in Supplemental Table 1.

After a fixed time of air sampling (Supplemental Table 1), the content of the filters was eluted with 1 mL HBsAg dilution buffer (HBSAGQ2 Dil HepB). The HBsAg concentration was then quantified using the Elecsys HBsAg II quant II assay on the cobas e 411 analyzer and were then standardized for comparable performance. The effective concentration of HBsAg was determined using the unprocessed signal results from the cobas e 801 analytic unit in conjunction with the corresponding calibration data. The limit of detection (LoD) and the limit of blank (LoB) for the Elecsys HBsAg II quant II assay were 0.05 IU/mL and 0.005 IU/mL, respectively.

Under routine laboratory conditions, swab testing was performed for all cobas e analyzers included in the panel, except the cobas e 801 analytical unit in combination with cobas pro integrated solution, to assess surface contamination due to aerosol formation (the swab testing locations are shown in Supplemental Fig. 2). To replicate a worst-case scenario, swab testing assessed the accumulation of HBsAg following 8-hours of full operation (the typical length of a shift for a laboratory operator) without any cleaning. Filter elution and subsequent quantification of HBsAg was performed as previously described.

### AEROSOL FORMATION IN AN END-TO-END LABORATORY WORKFLOW

Aerosol formation in an end-to-end laboratory workflow was assessed across four different test scenarios (Supplemental Table 2) that reflect critical processes and locations within this workflow (Figs. 1 and 2). In brief, the instruments assessed across the four test scenarios were the cobas connection modules (CCM, scenario 1; Roche Diagnostics International), manual output station 1 (MO1, scenario 2 and 4b; Hitachi, Tokyo, Japan), the cobas p 612 pre-analytical unit (scenario 3 and 4; Roche Diagnostics International), and the cobas p 501 post-analytical unit (scenario 4a; Roche Diagnostics International).

**Fig. 1.**
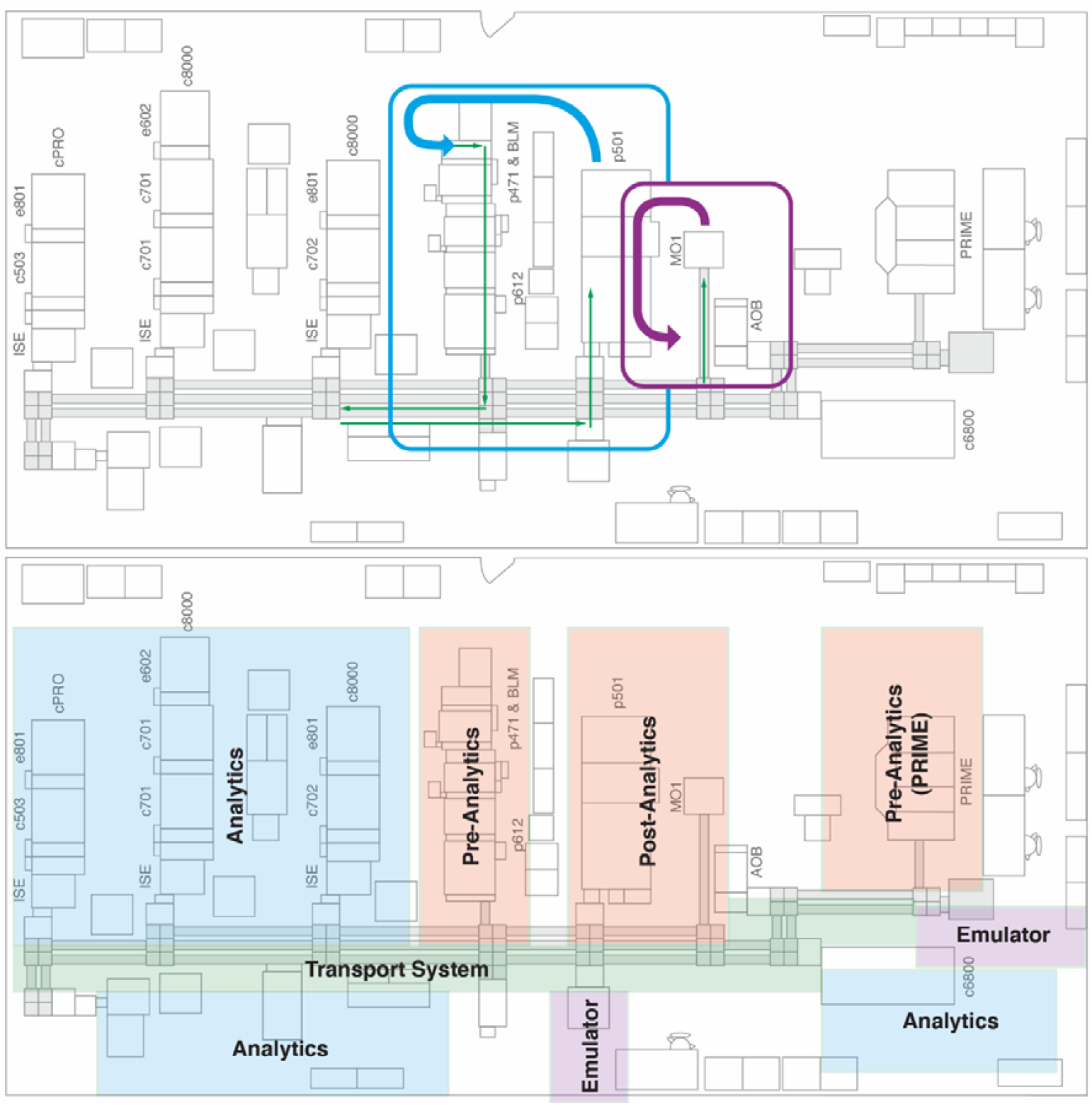
Floor plan of the end-to-end laboratory workflow sub-study. Arrows indicate the manual transport of sample tubes. Abbreviations: AOB, add-on buffer unit; BLM, bulk loader module; cPRO, cobas pro; c503, cobas 503 clinical chemistry analyzer; c701, cobas 701 clinical chemistry analyzer; c702, cobas 702 clinical chemistry analyzer; c6800, cobas 6800 molecular analyzer; c8000, cobas 8000 modular analyzer; e602, cobas 602 immune analyzer; e801, cobas e 801 analytical unit; ISE, ion selective electrode module; MO1, manual output station 1; PRIME, cobas PRIME, pre-analytical system; p471, cobas p 471 centrifuge unit; p501, cobas p 501 post-analytical system; p612, cobas p 612 pre-analytical unit.

**Fig. 2.**
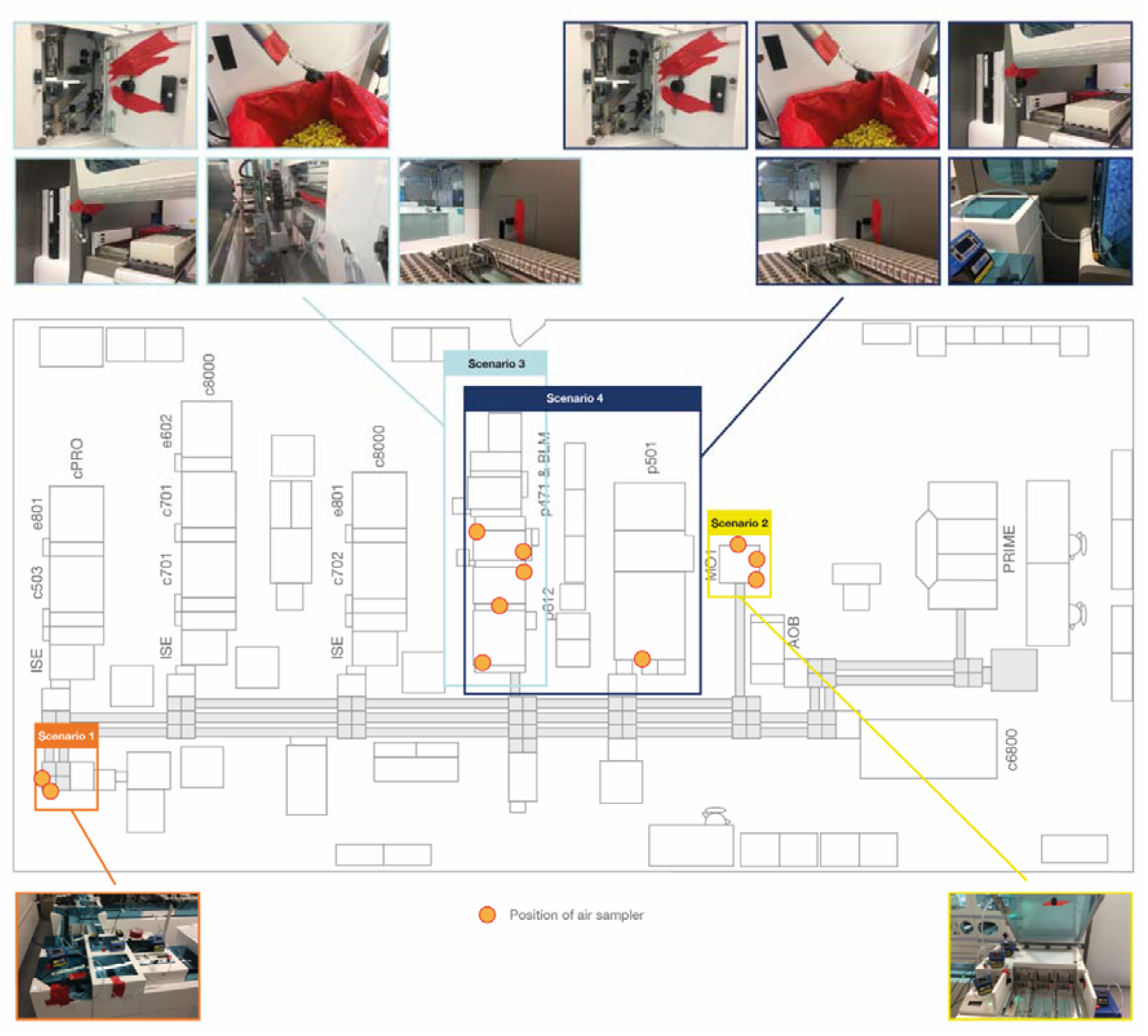
Placement of air sampling units in the end-to-end laboratory workflow. Abbreviations: AOB, add-on buffer unit; BLM, bulk loader module; cPRO, cobas pro; c503, cobas 503 clinical chemistry analyzer; c701, cobas 701 clinical chemistry analyzer; c702, cobas 702 clinical chemistry analyzer; c6800, cobas 6800 molecular analyzer; c8000, cobas 8000 modular analyzer; e602, cobas 602 immune analyzer; e801, cobas e 801 analytical unit; ISE, ion selective electrode module; MO1, manual output station 1; PRIME, cobas PRIME, pre-analytical system; p471, cobas p 471 centrifuge unit; p501, cobas p 501 post-analytical system; p612, cobas p 612 pre-analytical unit.

The test scenarios were chosen to allow assessment of aerosol formation during pre-analytical sample preparation, sample transport to the analyzers and automated output or archive solutions. Within the test scenarios, different sample tubes (Supplemental Table 2) were used to represent the different closure types and the most common sample tube dimensions. In scenarios 1 and 2, samples tubes were also selected for their relatively small filling volume, which allows a high filling level with a limited amount of HBsAg solution. A TeraTerm (TT) (Hitachi, Tokyo, Japan) was used in scenarios 1 and 2 to enable processing of 5-position racks directly on a linear conveyor without the use of the cobas p 612 pre-analytical unit as an entry point for the sample tube to the transport system. Similar to the cobas e analyzer panel sub-study, samples were processed using the pipetting sequences and application of the Elecsys SARS-CoV-2 Antigen test.

The air sampling pump and operating procedure previously described for the cobas e analyzer panel sub-study was also applied to this sub-study. Air sampling devices were positioned to measure aerosol formation at handling and transport locations (Fig. 2). Following air sampling, the elution and quantification of HBsAg concentration was performed in the same way as previously described for the cobas e analyzer panel sub-study but was conducted on the cobas e 801 analytical unit.

### APPLICATION TO A SARS-COV-2 CONTEXT

The quantified HBsAg values were applied to a SARS-CoV-2 context using the following assumptions: the typical air intake of a human is 600 L/h; the positive sample used in the experimental model represents a sample from a highly infectious SARS-CoV-2 individual with a viral load of 5×10^8^ viral RNA copies/mL (14); the effective dose needed to cause infection by SARS-CoV-2 is between 100–1000 viral RNA copies/mL (14).

For air sampling in both studies, the HBsAg uptake for an individual was calculated by dividing a typical human air intake of 600 L/h by the volume of air drawn through the air sampling filter and then multiplied by the HBsAg reduction factor (calculated by dividing the concentration of aerosol HBsAg particles measured during air sampling by the positive HBsAg sample concentration). The HBsAg uptake for an individual, in terms of viral RNA copies, was then calculated by multiplying the HBsAg uptake for an individual by the viral load (assumed to be 5×10^8^ viral RNA copies/mL). In the end-to-end laboratory workflow sub-study, as only samples containing HBsAg were used, the HBsAg uptake by an individual, in terms of viral RNA copies, was divided by 10 as it was assumed that 10% of the SARS-CoV-2 samples in a laboratory would be positive; therefore, mirroring the cobas e analyzer panel sub-study and replicating a worst-case real-world laboratory setting. An overview of the calculations used for air sampling in the study is provided in Table 1. For swab testing, the estimated maximum viral RNA copies were calculated by multiplying the HBsAg reduction factor (calculated in the same way as the air sampling data), by 5×10^8^ (representing a viral load of 5×10^8^ viral RNA copies/mL; Table 2).

**Table 1.**
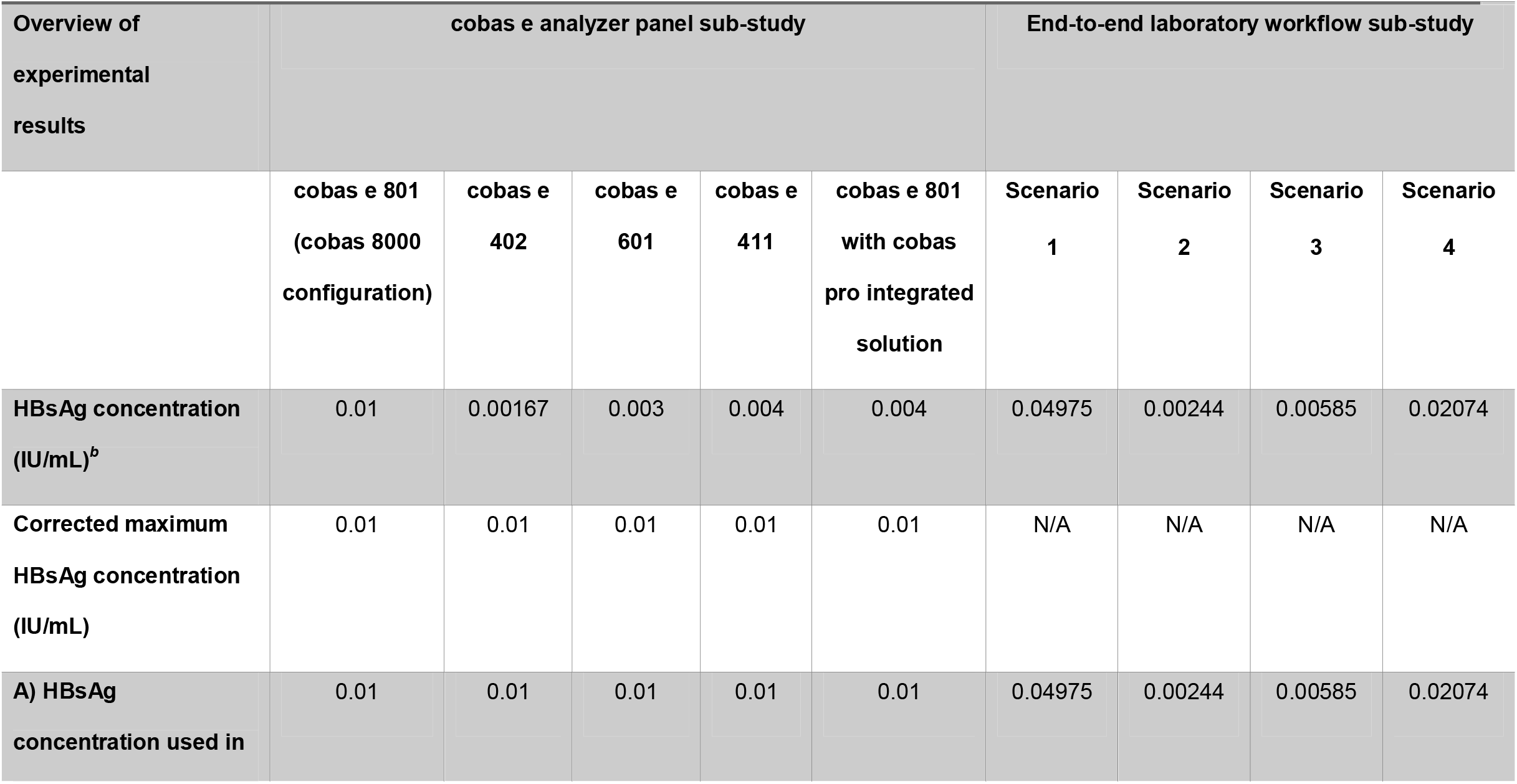
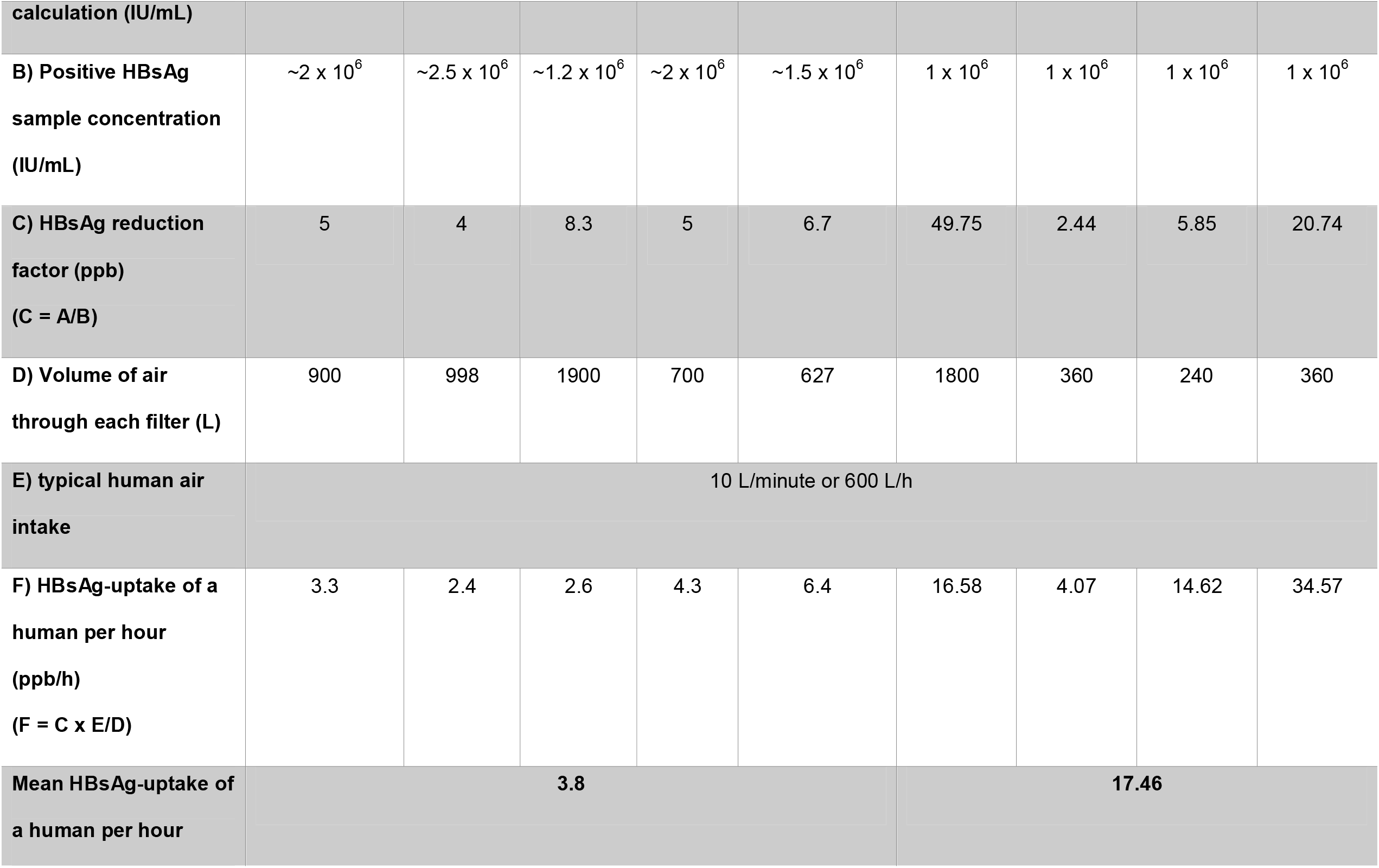
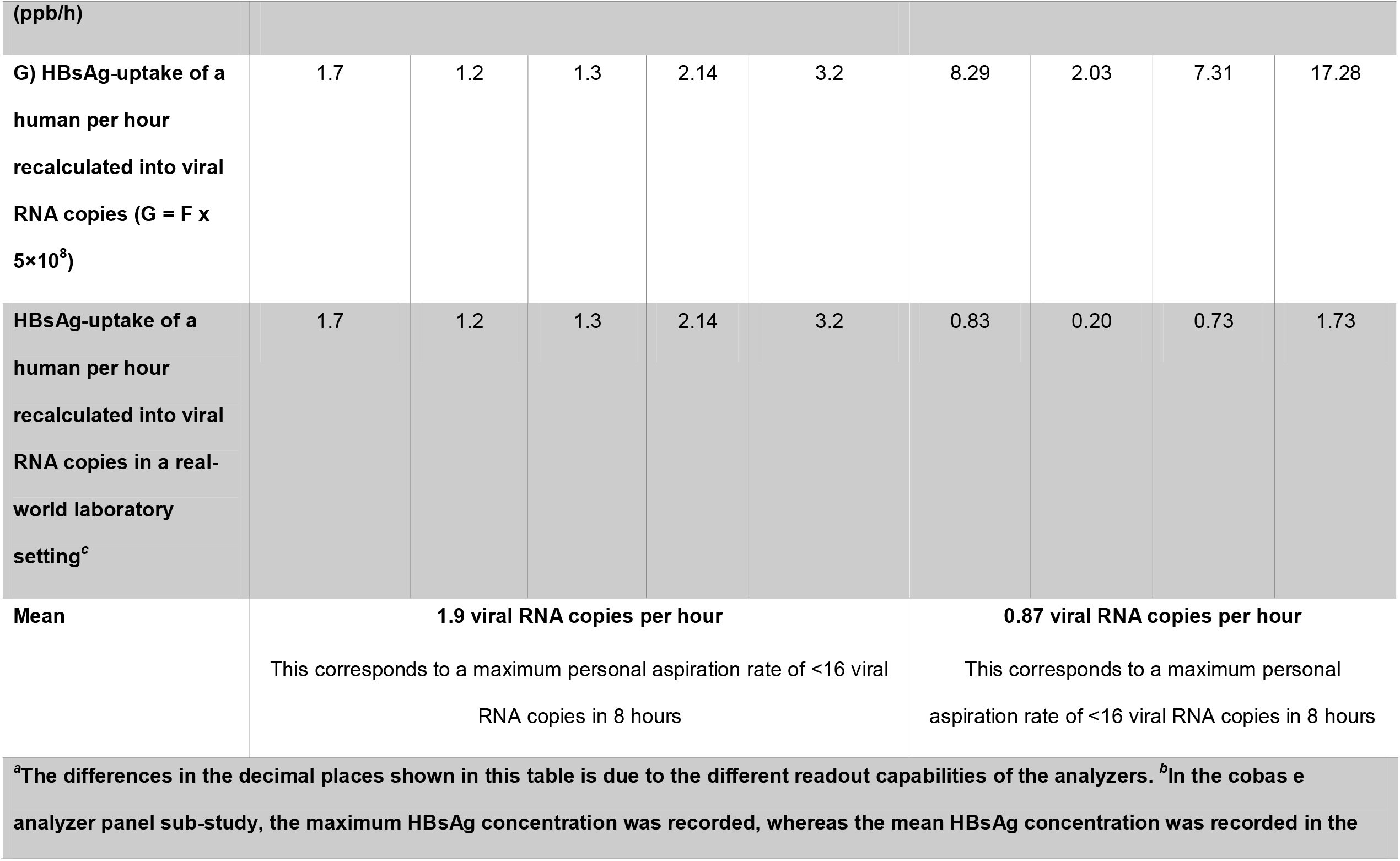
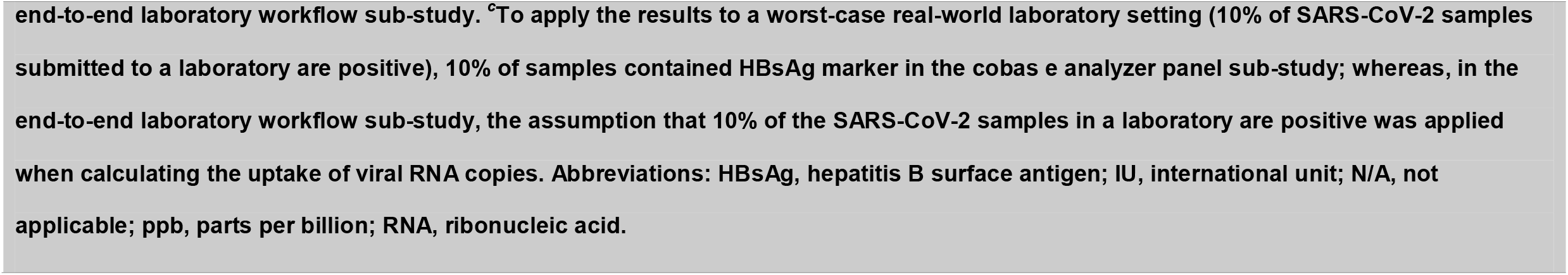
Overview of the calculations and results for air sampling^*a*^.

**Table 2.** Overview of the calculations and results for swab testing^*a,b*^.

| Overview of experimental results | cobas e analyzer panel sub-study |  |  |  |  |  |  |  |
| --- | --- | --- | --- | --- | --- | --- | --- | --- |
|  | cobas e 801 (cobas 8000 configuration)<sup>c</sup> |  | cobas e 402 |  | cobas e 601 |  | cobas e 411 |  |
|  | Before HBsAg assay run | After HBsAg assay run | Before HBsAg assay run | After HBsAg assay run | Before HBsAg assay run | After HBsAg assay run | Before HBsAg assay run | After HBsAg assay run |
| A) HBsAg concentration (IU/mL)<sup>d</sup> | - | 0.102 | 0.0024 | 0.0016 | 0.0024 | 0.022 | 0.004 | 0.004 |
| B) Positive HBsAg sample concentration (IU/mL) | ~2 x 10<sup>6</sup> | ~2 x 10<sup>6</sup> | ~2.5 x 10<sup>6</sup> | ~2.5 x 10<sup>6</sup> | ~1.2 x 10<sup>6</sup> | ~1.2 x 10<sup>6</sup> | ~2 x 10<sup>6</sup> | ~2 x 10<sup>6</sup> |
| C) HBsAg reduction factor (ppb) (C = A/B)<sup>d</sup> | - | 51.0 | 0.96 | 0.64 | 2.0 | 18.3 | 2.0 | 2.0 |
| Estimated maximum<br>viral RNA copies<br>(C x 5×10<sup>8</sup>)<sup>d</sup> | - | 25.5 | 0.48 | 0.32 | 1.0 | 9.2 | 1.0 | 1.0 |

<sup>a</sup>Swab testing assessed the accumulation of HBsAg over one day of full operation without any cleaning to replicate a worst-case scenario.
<sup>b</sup>The differences in the decimal places shown in this table is due to the different readout capabilities of the analyzers. <sup>c</sup>Data were not collected before the HBsAg assay run for the cobas e 801 analytical unit (cobas 8000 configuration). <sup>d</sup>For each cobas e analyzer, the values presented are the mean results from all different swab testing locations. Abbreviations: HBsAg, hepatitis B surface antigen; IU, international unit; ppb, parts per billion; RNA, ribonucleic acid.

### DATA PROCESSING AND ANALYSIS

Internally validated software (OASEpro HetIA, Roche Diagnostics International) was used to calculate the concentration (IU/mL) of HBsAg from the signal detected by the cobas e analyzers and data were processed using Microsoft Excel (Microsoft, Redmond, WA).

## Results

### COBAS E ANALYZER PANEL SUB-STUDY

Air sampling points were set up at various positions around the cobas e analyzers, as shown in Fig. 3. The maximum concentration of aerosol HBsAg particles was 0.01 IU/mL (recorded from a filter positioned around a fan outlet of the cobas e 801 analytical unit; Table 1). To improve the robustness of the study, 0.01 IU/mL was used as the maximum concentration of HBsAg particles for all cobas e analyzers when calculating the HBsAg reduction factor (Table 1). The cobas e 801 analytical unit in combination with cobas pro integrated solution recorded the highest HBsAg uptake per hour and the highest HBsAg uptake per hour when recalculated into viral RNA copies (6.4 ppb/h and 3.2 particles, respectively; Table 1). The cobas e 402 analytical unit recorded the lowest HBsAg uptake per hour and the lowest HBsAg uptake per hour when recalculated into viral RNA copies (2.4 ppb/h and 1.2 particles, respectively; Table 1). Across all cobas e analyzers, the mean HBsAg uptake per hour when recalculated into viral RNA copies was 1.9 particles, which results in a maximum aspiration rate of <16 viral RNA copies during a typical 8-hour shift for a laboratory operator (Table 1).

**Fig. 3.**
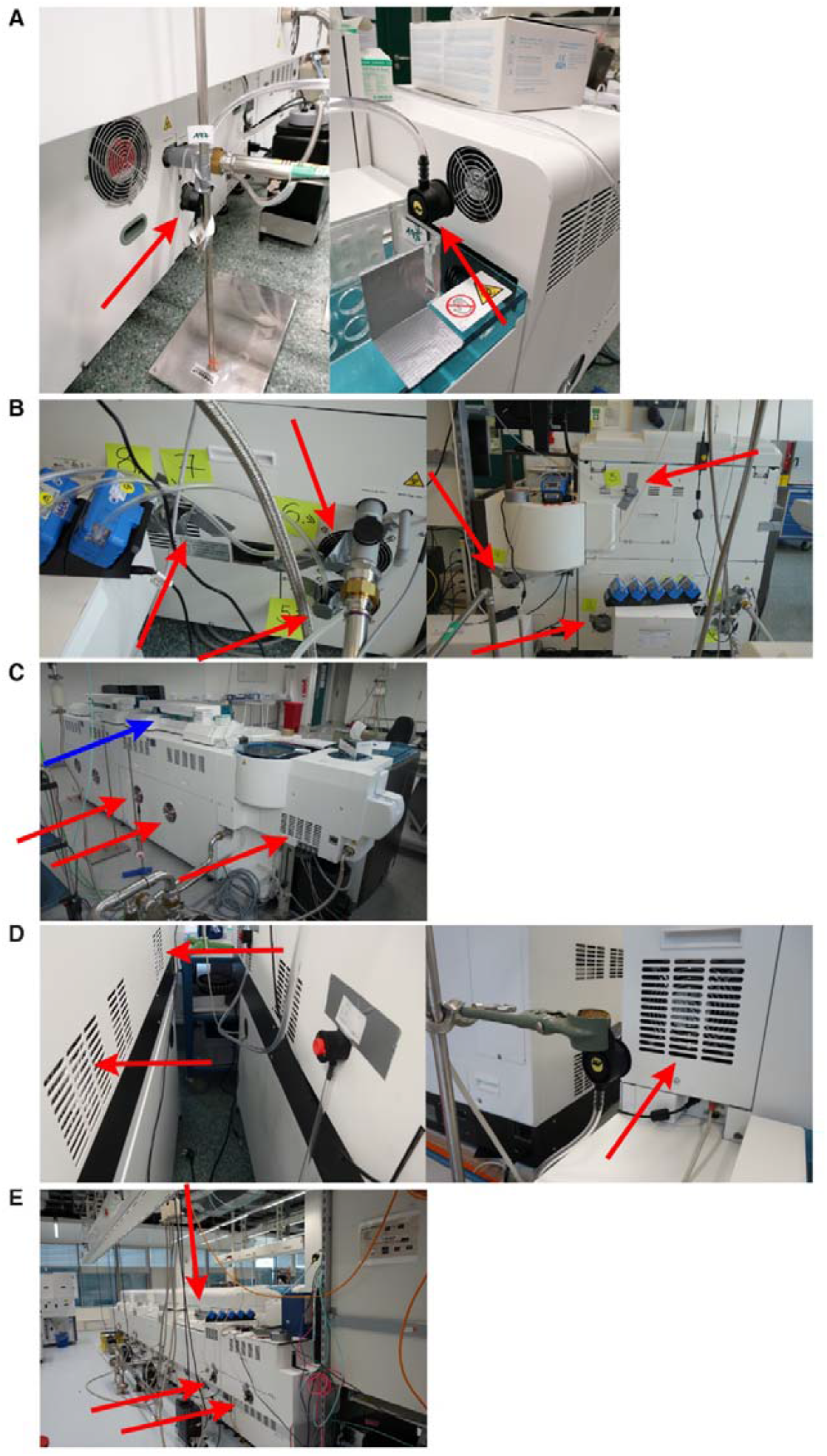
Selection of air sampling positions included in the cobas e analyzer panel sub-study^*a*^. (A) cobas e 801 analytical unit (cobas 8000 configuration). (B) cobas e 402 analytical unit (C) cobas e 601 module. (D) cobas e 411 analyzer. (E) cobas e 801 analytical unit in combination with cobas pro integrated solution. ^*a*^Positions of air sampling around fan outlets are indicated by red arrows and the sample pipette location is indicated by a blue arrow.

Swab testing around the cobas e 801 analytical unit, after the HBsAg assay run, recorded the highest mean estimated maximum viral RNA copies (25.5 particles; Table 2). Swab testing around the cobas e 402 analytical unit, after the HBsAg assay run, recorded the lowest mean estimated maximum viral RNA copies (0.32 particles; Table 2). For the cobas e 402 analytical unit, cobas e 601 module and cobas e 411 analyzers, there was minimal difference in the mean estimated maximum viral RNA copies before and after the HBsAg assay run (Table 2).

### END-TO-END LABORATORY WORKFLOW SUB-STUDY

Air sampling points were set up at various positions across the four test scenarios, as shown in Supplemental Fig. 3. In scenario 1 only, the eluate from the extracted filters exhibited an effective concentration of HBsAg particles above the LoD (0.05 IU/mL) of the Elecsys HBsAg II quant II assay (Fig. 4). For scenario 1, HBsAg levels above the LoD were also evident in the corresponding negative control and this result was not observed in the subsequent repetitions of the experiment (Fig. 4).

**Fig. 4.**
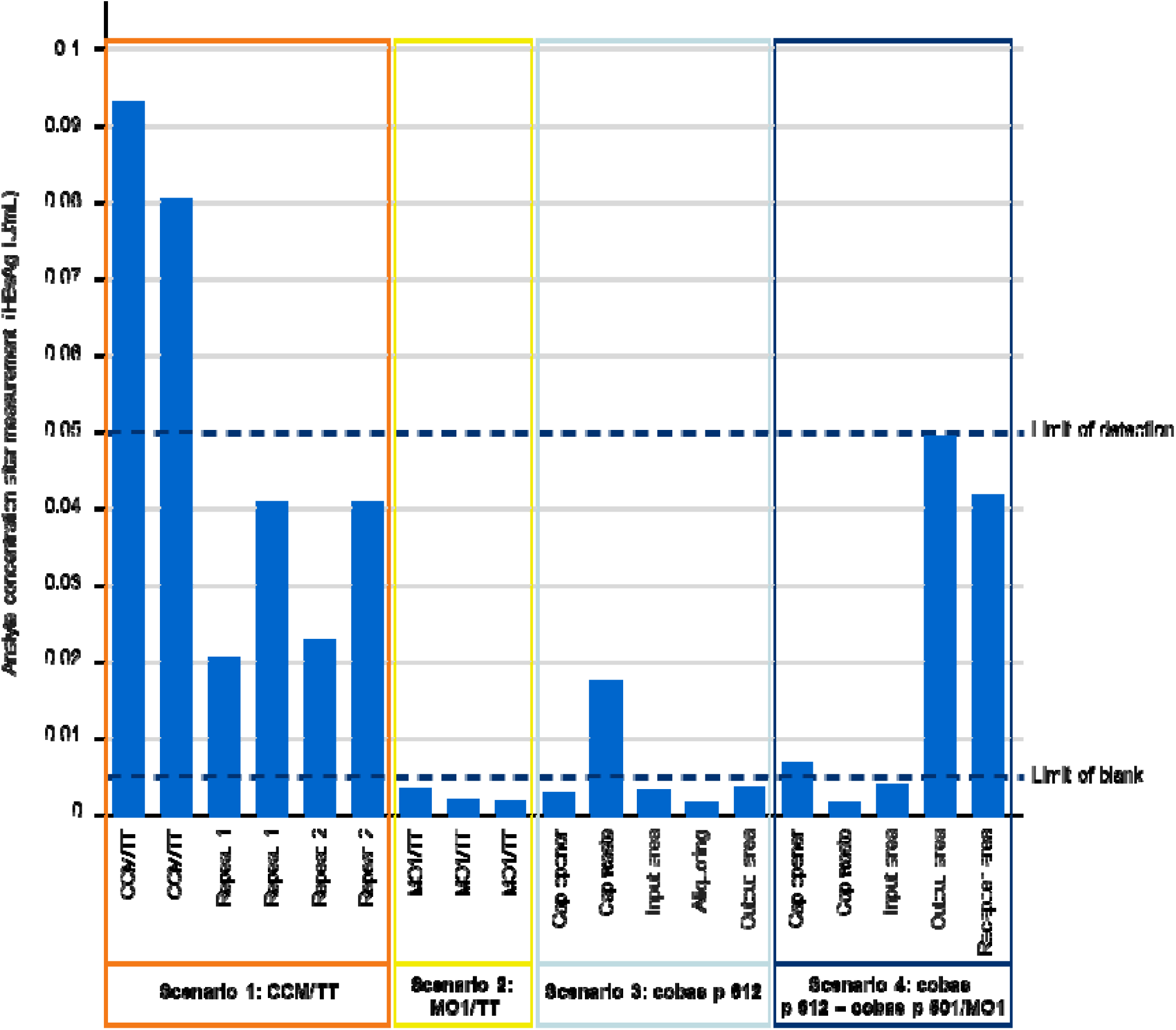
The effective HBsAg analyte concentration from each of the testing scenarios investigated in the end-to-end laboratory workflow sub-study. Quantification of HBsAg was performed on the cobas e 801 analytical unit. Abbreviations: CCM, cobas connection module; HBsAg, hepatitis B surface antigen; IU, international unit; MO1, manual output station 1; TT, TeraTerm.

Scenario 1 produced the highest mean concentration of aerosol HBsAg particles (0.04975 IU/mL) across the tested scenarios (Table 1). Scenarios 4 and 2 produced the highest and lowest HBsAg uptake per hour (34.57 ppb/h and 4.07 ppb/h, respectively; Table 1). With the assumption that 10% of the SARS-CoV-2 samples in a laboratory are positive, scenarios 4 and 2 also produced the highest and lowest HBsAg uptake per hour when recalculated into viral RNA copies (1.73 particles and 0.20 particles, respectively; Table 1), while the corresponding mean across the test scenarios was 0.87 particles (Table 1). As such, there is a mean maximum aspiration rate of <16 viral RNA copies during an 8-hour shift.

## Discussion

To our knowledge, this is the first study to assess formation of potentially infectious aerosols when processing infectious samples using a panel of cobas e analyzers and in an automated end-to-end laboratory workflow. While viral agents transmitted through blood and bodily fluids are the most common source of infection for personnel within a diagnostic laboratory (15), it is timely to assess aerosol infection risk given the present COVID-19 pandemic. In this study, data on potentially infectious aerosol formation were applied to a SARS-CoV-2 context and the low number of viral RNA copies detected inferred a minimal risk of SARS-CoV-2 infection to laboratory operators when processing potentially infectious samples.

In the end-to-end laboratory workflow sub-study, in scenario 1 (the CCM/TT units), elevated levels of HBsAg above the LoD were evident, elevated levels were also evident in the corresponding negative control; however, such levels were not present in the subsequent repetitions of the experiment. This can be considered a contamination-driven effect and was not considered to originate from the samples processed on the CCM/TT units.

Hepatitis B is spread through contact with infected bodily fluids; however, HBsAg can self-assemble to virus-like particles that are assumed to distribute via aerosols; therefore, the use of recombinant HBsAg in the model system allows the model to be applied to all airborne pathogens and not just SARS-CoV-2. This addresses an important unmet clinical need as aerosol formation has previously been undervalued in terms of transmission of respiratory viral diseases due to a lack of understanding of how infectious aerosols are produced and transported (16).

In this study, the mean maximum aspiration rate detected was <16 viral RNA copies during an 8-hour shift when using cobas e analyzers and in an end-to-end laboratory context. Furthermore, for swab testing, the greatest number of estimated maximum viral RNA copies was limited to 25.5 particles over 8-hours of full operation and there was only a minimal difference in mean estimated maximum viral RNA copies before and after the HBsAg assay run. The infectious dose of a virus can vary greatly among respiratory viruses (17). For SARS-CoV-2, the minimum dose of virus particles necessary to cause infection remains an active research question; however, it has been estimated to be between 100 and 1000 particles (18-21). Therefore, the risk of SARS-CoV-2 infection for laboratory operators when using cobas e analyzers and in an end-to-end laboratory workflow can be considered remote. Applying the outcomes of studies of other viruses that have evaluated risk of infection to laboratory operators may not be relevant in a SARS-CoV-2 context due to the highly infectious nature of SARS-CoV-2; generally inhaled viruses require 1950–3000 particles to cause infection, compared with between 100 and 1000 particles for SARS-CoV-2 (18-21).

When applying the findings of the study to a SARS-CoV-2 context, the assumption regarding viral load did not consider the effect of variants of SARS-CoV-2. For example, in the first known community transmission event of the delta variant in mainland China, it was shown that the viral load associated with the delta variant was approximately 1000 times higher than with the alpha or beta strain (22). In addition, vaccination status has also been shown to affect viral load kinetics, whereby vaccinated individuals have a faster mean rate of viral load decline relative to unvaccinated individuals (23). Some European countries have already made vaccination against COVID-19 mandatory for all healthcare workers (24, 25).

A strength of this study is that a worst-case scenario (10% of samples were highly positive) was tested in several different analyzers using a clinically relevant end-to-end laboratory setup; thus, replicating a real-world testing laboratory for SARS-CoV-2. A further strength of this study is that swab testing of the instrument surfaces was performed, which, in addition to aerosol formation, could represent a source of infection for laboratory workers (26-29). In a similar study, Farnsworth et al. used a fluorescent marker to assess instrument and specimen contamination and found that while there is a low risk of instrument contamination, the handling of infectious specimen containers can contaminate laboratory surfaces (30). Relative to a fluorescent marker, the use of the HBsAg model in this study was more representative of the mass: volume ratio seen in the aerosol distribution of virus particles. Correct adherence in the use of personal protective equipment (PPE) can be very effective in preventing infection through contact with contaminated surfaces (30) and inhalation of droplets associated with SARS-CoV-2 infection (31, 32). Van Doremalen et al. found that under experimental conditions, SARS-CoV-2 is viable in aerosols for 3 hours (9); therefore, PPE should always be worn in a laboratory setting. Healthcare workers caring for patients with COVID-19 report adherence rates of 98.6% to PPE protocols (33); similar adherence rates are important to reduce SARS-CoV-2 infection risk in a laboratory context. Future work should assess molecular modules, to expand upon the findings for analyzers shown here.

In conclusion, the low production of potentially infectious aerosols when using cobas e analyzers and in an end-to-end laboratory workflow (which includes processes of pre-analytical sample preparation, sample transport to analyzers and automated output or archive solutions) is consistent with a remote risk of SARS-CoV-2 infection for laboratory operators. These results can be considered reassuring to laboratory operators involved in the processing of potentially infectious samples.

## Data Availability

The data supporting this manuscript can be requested in writing from the corresponding author.

## Ethical Guidelines

The artificial samples used for this study were produced using purified, vendor purchased hepatitis B surface antigen (HBsAg) protein and dilution media; therefore, no ethical approval was needed.

## Author Contributions

All authors confirmed they have contributed to the intellectual content of this paper and have met the following 4 requirements: (a) significant contributions to the conception and design, acquisition of data, or analysis and interpretation of data; (b) drafting or revising the article for intellectual content; (c) final approval of the published article; and (d) agreement to be accountable for all aspects of the article thus ensuring that questions related to the accuracy or integrity of any part of the article are appropriately investigated and resolved.

## Authors’ Disclosures or Potential Conflicts of Interest

### Employment or Leadership

Markus Eckl, Doris Huether, Oliver Larbolette, Alessia Lo Faso, Beatus Ofenloch-Haehnle, Marlene Riesch and Rolf Herb are employed by Roche Diagnostics GmbH. Géza Burghardt is employed by Roche Diagnostics International Ltd.

### Consultant or Advisory Role

Markus Eckl participated in the Roche Diagnostics SARS-CoV-2 Biosafety, Antigen Testing & Infectivity Advisory Board Meeting (8^th^ December 2020)

### Stock Ownership

Géza Burghardt, Markus Eckl, Oliver Larbolette, Beatus Ofenloch-Haehnle and Rolf Herb hold non-voting equities in F. Hoffman-La Roche

### Honoraria

N/A

### Research Funding

Roche Diagnostics International Ltd funded the study and third-party medical writing assistance

### Expert Testimony

N/A

### Patents

N/A

### Other Renumeration

N/A

## Acknowledgements

The authors wish to thank Marie Christine Weiss, PhD, for providing additional support in planning/reviewing experimental concepts, Thomas Grau, PhD, (as part of Portfolio Solution Integration at Roche Diagnostics International Ltd) for financing of the end-to-end laboratory workflow sub-study and Ursula Giesen, PhD, for a comprehensive review of both the work conducted in the cobas e analyzer panel sub-study and subsequent reports. Third-party medical writing support for the development of this manuscript, under the direction of the authors, was provided by Luke Edmonds, MSc, and Heather Small, PhD, of Ashfield MedComms (Macclesfield, UK), an Ashfield Health company, and was funded by Roche Diagnostics International Ltd (Rotkreuz, Switzerland). COBAS, COBAS E, COBAS P, COBAS PRIME, COBAS PRO and ELECSYS are trademarks of Roche. All other product names and trademarks are property of their respective owners.

## Abbreviations (in order of mention)

COVID-19: coronavirus disease 2019
SARS-CoV-2: severe acute respiratory syndrome coronavirus 2
HBsAg: hepatitis B surface antigen
LoD: limit of detection
LoB: limit of blank
CCM: cobas connection modules
MO1: manual output station 1
TT: TeraTerm
PPE: personal protective equipment

