## Supplementary materials for "Aerosol Formation During Processing of Potentially Infectious Samples on Roche Immunochemistry Analyzers (cobas e analyzers) and in an End-to-End Laboratory Workflow to Model SARS-CoV-2 Infection Risk for Laboratory Operators"

**Supplemental Material**

**SUPPLEMENTAL METHODS**

**Air Sampling Plausibility Checks in the cobas e Analyzer Panel Sub-Study**

In the first air sampling plausibility check, a highly positive hepatitis B surface antigen (HBsAg) sample (2 million IU/mL) was subject to mechanical agitation (stirring) and multiple pipetting between an open beaker and vial placed in a bio-safety cabinet (approximately 1 m³) without air flow for 15 minutes. The air sampling device, Gilian GilAir Plus Air Sampling Pump (Sensidyne, St. Petersburg, Florida), was set up next to the beaker as shown in Supplemental Figure 1, A and B at 2000 mL/min. After 90 minutes and 179 L of air filtered, the polycarbonate filter (0.8 μm pore size; SKC Inc., Omega Division, Eighty Four, Pennsylvania) was eluted with HBsAg-negative elution buffer (HBSAGQ2 Dil HepB, Roche Diagnostics International, Rotkreuz, Switzerland) for quantitative determination.

In the second air sample plausibility check, a highly positive HBsAg sample was pipetted between two sample cups placed in a bio-safety cabinet (approximately 1 m³) without air flow for 15 minutes (Supplemental Figure 1C). The air sampling device, Gilian GilAir Plus Air Sampling Pump, was set up next to the beaker as shown in Supplemental Figure 1A. The first cup contained 1 mL of highly positive HBsAg sample and the second cup contained 200 μL of the same sample; 200 μL of the highly positive HBsAg sample was pipetted between cups every 30 seconds, for a total of 15 minutes as shown in Supplemental Figure 1D. After this, the polycarbonate filter (0.8 μm pore size) was eluted with HBsAg-negative elution buffer (HBSAGQ2 Dil HepB) for quantitative determination. The following variables were tested in this setup: distance of the air sampler from the sample pipetting point (4 versus 20 cm), air flow ie, pump speed of the filter pump unit (2000 versus 3000 mL/min), and the pore size of the filter (0.4 versus 0.8 μm).

**Supplemental Tables**

| Supplemental Table 1. Overview of the Experimental Conditions For Air Sampling in the cobas e Immunoanalyzer Panel Sub-Study | | | | | |
| --- | --- | --- | --- | --- | --- |
| Overview on Conditions in Experiments | **cobas e 801 (cobas 8000 Configuration)** | **cobas e 402** | **cobas e 601** | **cobas e 411** | **cobas e 801 With cobas pro Integrated Solution** |
| Laboratory settings |  |  |  |  |  |
| air volume  (m³) | 300 | 300 | 300 | 150 | Approx. 430 |
| air exchange  rate   (per hour) | 3 | 3 | 3 | 3 | 2.5 |
| total air  volume  exchanged  (m³/h) | 900 | 900 | 900 | 450 | 1075 |
| Positive HBsAg sample concentration (IU/mL) | ~2 x 10^6^ | ~2.5 x 10^6^ | ~1.2 x 10^6^ | ~2 x 10^6^ | ~1.5 x 10^6^ |
| Ratio positive:negative samples | 1:9 | 1:9 | 1:9 | 1:9 | 1:3 |
| Number of repeat experiments | 3 | 2 | 2 | 2 | 2 |
| Number of air collection filters | 6 | 9 | 7 | 5 | 5 |
| Supplemental Table 1. Continued | | | | | |
| Distribution of air collection filters | - Around fan outlets (x3)  - Standard laboratory operator position (x2)  - Laboratory room (x1) | - Around fan outlets (x7)  - Standard laboratory operator position (x1)  - Computer workstation (x1) | - Around fan outlets (x4)  - Standard laboratory operator position (x2)  - Computer workstation (x1) | - Around fan outlets (x3)  - Standard laboratory operator position (x1)  - Laboratory room (x1) | - Around fan outlets (x3)  - Standard laboratory operator position (x1)  - Computer workstation (x1) |
| Air sampling time (h) | 8 | 8.3 | 8 | 5.8 | 4.5 |
| Volume of air passed through each filter (L) | 900 | 998 | 1900 | 700 | 627 |

Abbreviations: IU, international unit.

| Supplemental Table 2. Different Test Scenarios Investigated In The End-To-End Laboratory Workflow Sub-Study | | | | | |
| --- | --- | --- | --- | --- | --- |
| Scenario No. | **Setup** | **No. of Samples** | **Description** | **Rationale for Setup** | **Filter Positioning** |
| 1 | CCM/TT | 4 | - Two 5-position racks were loaded with one or two 2.5 mL mini sample cups and contained 2 mL undiluted marker solution - Sample cups were processed in an ‘endless’ loop overnight - Overnight aerosol capturing procedure was repeated three times - Extraction of filter was completed with the HBsAg dilution solution provided with the corresponding reagent kit - Sample tubes were then processed on the cobas e 801 analytical unit - Total air sampling time was ~15 hours | - Aerosol formation is assessed during sample transport of open tubes on the CCM | - Filters were positioned inside the TT, inside the analytical connector 05 |
| Supplemental Table 2. Continued | | | | | |
| 2 | MO1/TT | 900 | - 15 5-position racks were loaded with either 0.5 or 1 mL marker solution, or both in either 1 mL micro or 2.5 mL mini sample cups, respectively - The scenario was repeated 30 times to cover each position of the uppermost level of MO1 output area - Total air sampling time was ~3 hours | - Allows assessment of aerosol formation during processing of post-analytical samples (automated output or archive solutions) - Reflects the transport of open tubes and loading on the MO1 output station | - Three filters were positioned inside the MO1 |
| Supplemental Table 2. Continued | | | | | |
| 3 | cobas p 612 | 15 | - Five sample tubes (screw cap Greiner VACUETTE only) were prefilled with 6.5 mL marker solution - Sample tubes were loaded on the BLM and centrifuged for 60 seconds at 3900 rpm - Sample tubes with maximum filling volume were used to produce a high amount of aliquots - No CCM transport system was used (sample tubes were placed on the output area of cobas p 612) - Following aliquoting, the secondary sample tubes produced were pooled back into the original primary sample tube and then reloaded on the BLM to repeat the workflow - Overall, the experiment was repeated three times and 150 aliquots were generated - Total air sampling time was ~4 hours | - Allows assessment of aerosol formation during the processing of pre-analytical samples - Contains the loading on the BLM, centrifugation, decapping and aliquoting. | - Filters were positioned inside the decapper area; at the tube cap waste area; at the cobas p 612 input area; near the aliquoting area; and at the cobas p 612 output area |
| Supplemental Table 2. Continued | | | | | |
| 4a | end-to-end: cobas p 612 – cobas p 501 | 45 | - 15 sample tubes (Greiner VACUETTE and Becton-Dickinson tubes) filled with marker solution were loaded to cobas p 612 and then transported to cobas p 501 - Sample tubes were retrieved from cobas p 501 then manually decapped (if needed) - Sample tubes were recapped and reloaded on cobas p 612 - 30 sample tubes that were compatible with cobas p 471 underwent centrifugation (this represents a worst-case scenario for the non-centrifuged tubes because of a potentially higher risk of aerosol formation, caused by remaining sample fluid on the upper inner part of the tube and/or tube cap) - This was repeated three times - Total air sampling time ~3 hours | - Allows assessment of aerosol formation during the processing of pre-analytical samples and post-analytical samples (automated output or archive solutions) - Shows the end-to-end process of centrifuged and non-centrifuged samples (for example, decapping without aliquoting on cobas p 612, followed by transport via the CCM to cobas p 501 and MO1, respectively) | - Filters were positioned inside the decapper area; at the tube cap waste area; at the cobas p 612 input area; at the cobas p 612 output area; and connected to CCM, near the recapping area |
| 4b | end-to-end: cobas p 612 – MO1 | 15 | - Five sample tubes (Roche cobas PCR media tubes) filled with marker solution were loaded to cobas p 612 and then transported to MO1 - Sample tubes were retrieved from MO1 then manually decapped (if needed) - Sample tubes were recapped and reloaded on cobas p 612 - No sample tubes underwent centrifugation - This was repeated three times - Total air sampling time ~3 hours |  |  |

Abbreviations: BLM, bulk loader module; CCM, cobas connection modules; HBsAg, hepatitis B surface antigen; MO1, manual output station 1; PCR, polymerase chain reaction; TT, TeraTerm.

**Supplemental Figures**

**Supplemental Figure 1.** *Air sampling plausibility checks.* *(A–B) Gilian GilAir Plus Air Sampling Pump next to the beaker, used in both air sampling plausibility checks. (C) overview of the setup in the second air sampling plausibility check. (D) manual pipetting in the second air sampling plausibility check.*

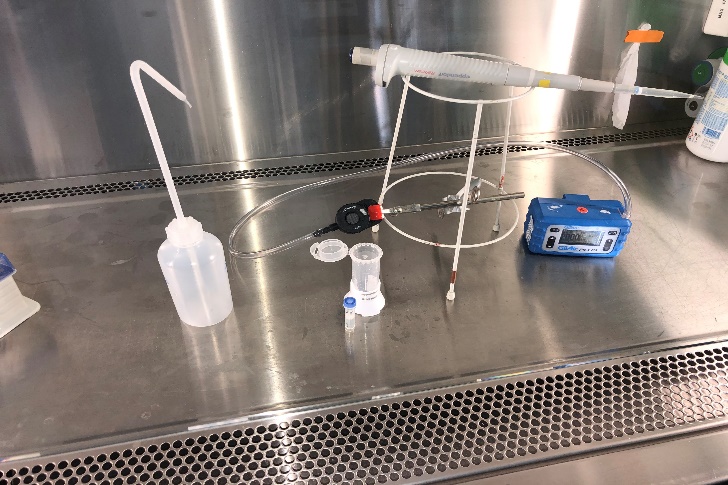

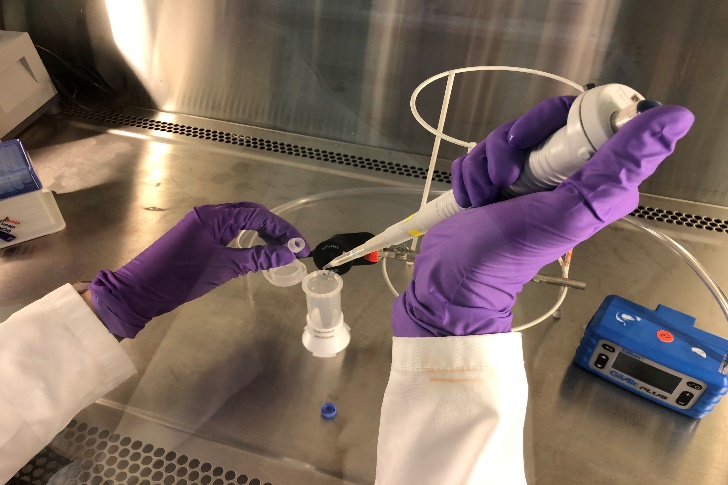

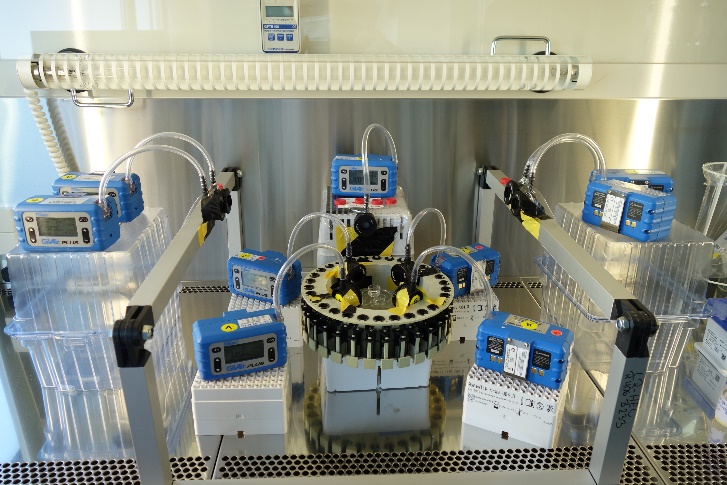

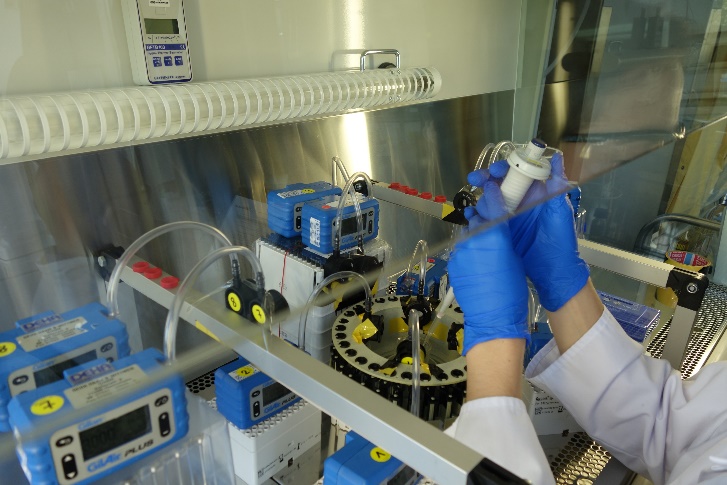

**A**

**B**

**C**

**D**

**Supplemental Figure 2.** *Swab testing locations.* *Swab testing was performed around the cobas e 801 analytical unit (cobas 8000 configuration) (A)^a^; cobas e 402 analytical unit (B)^b^; cobas e 601 module (C)^b^; and cobas e 411 analyzer (D)^c^.*

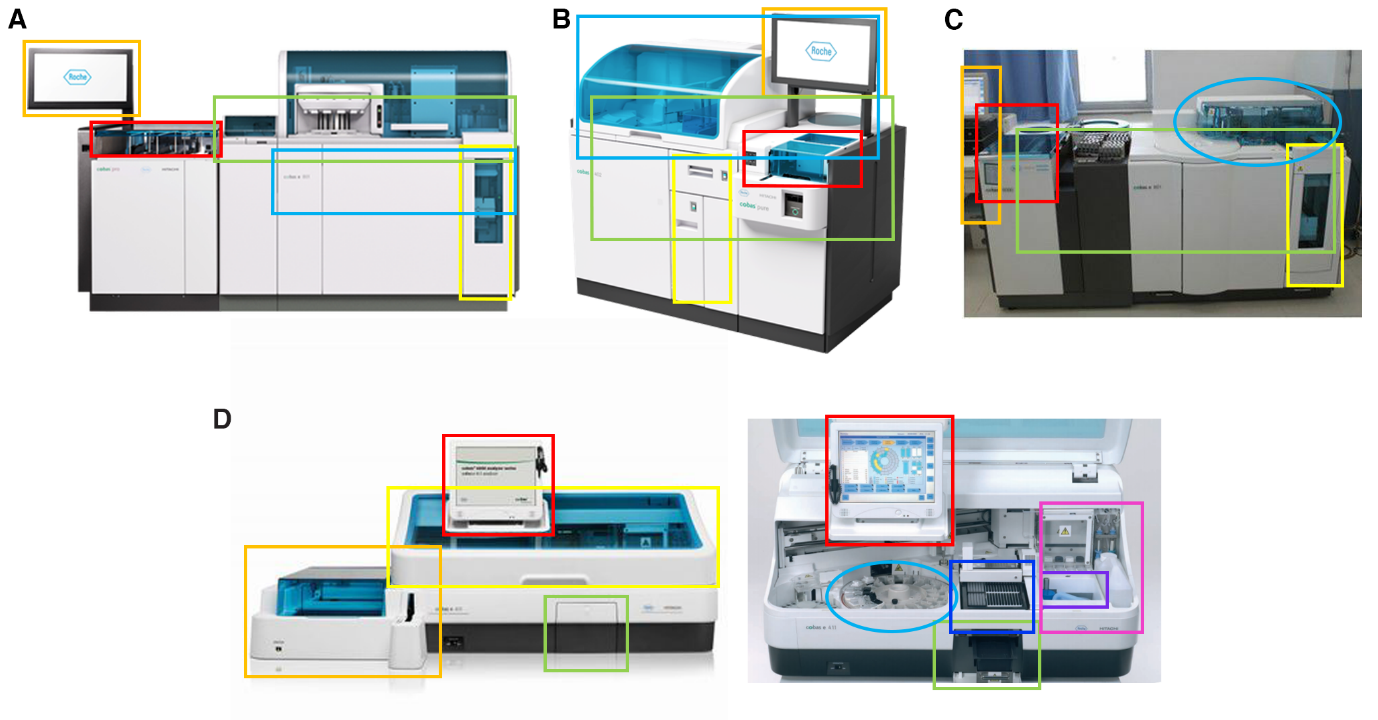

*^a^ Swab testing was performed at the computer keyboard, mouse and monitor (orange rectangle); at the core sample inlet/outlet and corresponding handles (red rectangle); at the front of the analytical unit, including the corresponding handles and surfaces (green rectangle); at the top cover of the analytical unit and corresponding surfaces (blue rectangle) at the waste drawer and surrounding areas, including handles (yellow rectangle).
^b^ Swab testing was performed at the upper surface of the immunoanalyzer (B, blue rectangle; C, blue oval); at the front of the immunoanalyzer, including handles (green rectangle); at the waste drawer (yellow rectangle); at the entrance for the samples and the corresponding surfaces (red rectangle); at the computer keyboard, mouse and monitor (orange rectangle).
^c^ Swab testing was performed at the sample rack (orange rectangle); at the hood of the immunoanalyzer (yellow rectangle); at the touch screen (red rectangle); at the solid waste area (green rectangle); at the reagent rotor (light blue oval); at the disposable waste area (dark blue rectangle); at the liquid waste area (purple rectangle); at the ProCell, CleanCell and SysWash areas (pink rectangle).*

**Supplemental Figure 3.** *Test scenarios used in the end-to-end laboratory workflow sub-study^a^. (A) Scenario 1 (CCM/TT). (B) Scenario 2 (MO1/TT). (C)^b^ Scenario 3 (cobas p* *612 pre-analytical unit). (D)^c^ Scenario 4 (4a, end-to-end: cobas p* *612 pre-analytical unit – cobas p* *501 post-analytical unit; 4b end-to-end: cobas p* *612 pre-analytical unit– MO1).*

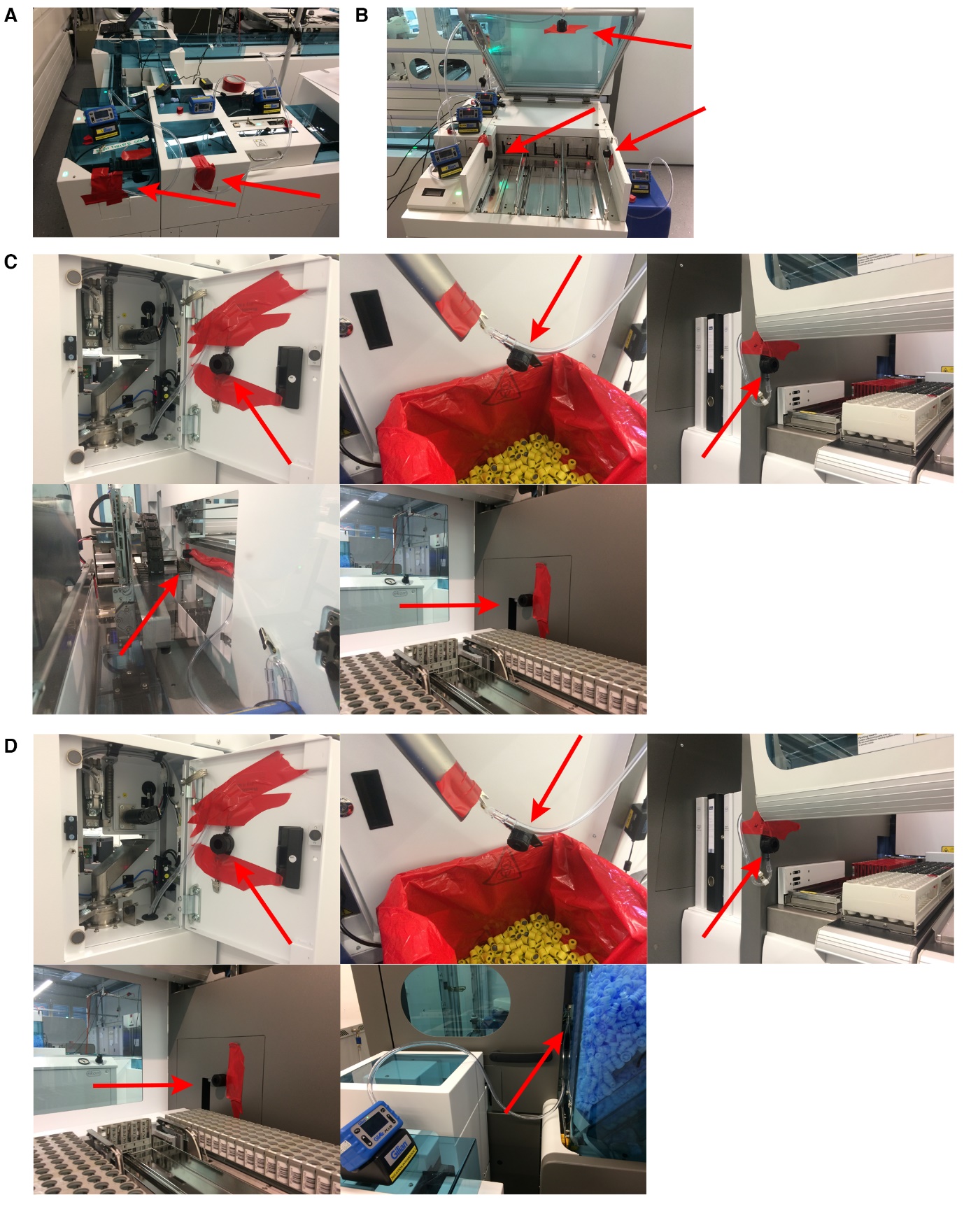

*Abbreviations: CCM, cobas connection module; MO1, manual output station 1; TT, TeraTerm. ^a^ Air filters are indicated by red arrows.
^b^ Pictured left to right: inside the decapper area; at the tube cap waste area; at the cobas p* *612 pre-analytical unit input area; near aliquoting area; and at the cobas p* *612 pre-analytical unit output area.
^c^ Pictured left to right:* *inside the decapper area; at the tube cap waste area; at the cobas p* *612 input area; at the cobas p* *612 pre-analytical unit output area; and connected to the cobas p* *501 post-analytical unit recapping area.*
